# An aortic hemodynamic fingerprint reduced order modeling analysis reveals traits associated with vascular disease in a medical biobank

**DOI:** 10.1101/2024.04.19.590260

**Authors:** Ryan Sokolow, Georgios Kissas, Cameron Beeche, Sophia Swago, Elizabeth W. Thompson, Mukund Viswanadha, Julio Chirinos, Scott Damrauer, Paris Perdakaris, Daniel J. Rader, Walter R. Witschey

## Abstract

**Purpose:** To determine the clinical relevance of reduced order model (ROM) aortic hemodynamic imaging-derived phenotypes (IDPs) for a range of flow conditions applied to computed tomography (CT) scan data in the Penn Medicine Biobank (PMBB).

**Methods:** The human thoracic aorta was automatically segmented in 3,204 chest CT scans from patients in the Penn Medicine Biobank (PMBB) patients using deep learning. Thoracic aorta anatomic IDPs such as aortic diameter and length were computed. Resistance, and flow boundary conditions, were varied, resulting in 125,000 ROM simulations, producing a fingerprint of aortic hemodynamics IDPs for a range of flow conditions. To determine the clinical relevance of the aortic hemodynamic fingerprint, untargeted phenome wide association studies (PheWAS) for disease conditions were performed using aortic geometries and pulse pressure as IDPs.

**Results:** By utilizing patient metadata from the PMBB, the human aortic radius for different age groups over a normalized radius was visualized, showing how the vessel deforms with age, as well as other characteristic geometric information. The average radius of the ascending thoracic aortic data set was 26.6 ± 3.1 mm, with an average length of 310 ± 37 mm. A combination of pathology codes (phecodes) and hemodynamic simulations were utilized to develop a relationship between them, showing a strong relationship between the resulting pulse pressure and diseases relating to aortic aneurysms and heart valve disorders. The average pulse pressure calculated by the model was 22.5 ± 8.5 mmHg, with the maximum pressure modeled by the system being 201 mmHg, with the minimum being 63.6 mmHg. The pulse pressures of the most significant phecodes were examined for patients with and without the condition, showing a slight separation between the two cases. The pulse pressure was also slightly negatively correlated with the calculated tapering angle of the ascending thoracic aorta.

**Conclusions:** ROM hemodynamic simulations can be applied to aortic imaging traits from thoracic imaging data in a medical biobank. The derived hemodynamic fingerprint, describing the response of the aorta to a range of flow conditions, shows clinically relevant associations with disease.

## Introduction

The aorta is the main delivery conduit for oxygenated blood from the left ventricle to the rest of the body. Aortic diseases can affect how well the blood is distributed, thus affecting the oxygenation of tissues all throughout the body. Aortic diseases affect at least 5.7% of individuals, the more common diseases being aortic dissection, aortic aneurysm, and atherosclerotic plaque [1, 2]. Ascending Thoracic Aortic Aneurysms (ATAAs) are a widening of the aorta due to the weakening of the aortic wall [3]. There is evidence that variations in hemodynamic properties such as flow velocity and pressure contribute to the development of ATTAs [3, 4]. Therefore, the ability to quickly model hemodynamic properties of the aorta for any given patient could potentially allow for clinical predictions on the progression of aortic diseases.

Longitudinal population-based studies have been used to determine the progression of aortic disease. The Multi-Ethnic Study of Atherosclerosis (MESA) is a prospective epidemiologic study of the incidence, risk factors and progression of atherosclerosis. A subset of these participants had CT imaging of abdominal aortic calcium in addition to coronary calcium, ventricular mass and function, flow-mediated brachial artery endothelial vasodilation, carotid intimal-medial wall thickness and distensibility of the carotid arteries [5]. MESA examined the role arterial pressure wave reflections played as a risk factor for critical heart failure, and discovered patients with a larger reflected wave magnitude were at a higher risk for heart failure [6]. Another study with similar aims was the Framingham Heart Study (FHS), which consisted of 1230 subjects with the goal to determine whether left ventricular mass is associated with an increased risk for stroke [7]. The subjects in the FHS are examined with echocardiography at 8-year intervals to measure left ventricular mass, as well as blood pressure, cholesterol levels and left atrial size.

Hemodynamic measurements in the aorta could provide additional information about subclinical disease preceding the development of ATAAs. While FHS and MESA measured the progression of atherosclerosis and other important geometric quantities for aortic disease, technological limitations prevented these studies from directly assessing high-resolution hemodynamic characteristics of the aorta (pressure and volumetric flow measurements). Such measurements could be obtained using invasive catheterization procedures. Alternatively, given appropriate boundary conditions, they could be obtained using computational fluid dynamics (CFD), a tool that utilizes computers and the governing equations of fluid motion to perform calculations. CFD could be used to obtain hemodynamic properties such as wall shear stress (WSS), pressure and velocity waveforms that would be clinically useful in understanding the development of aortic pathologies [8, 9].

While CFD can provide useful information on the hemodynamic properties of the aorta, there are limitations to its usefulness in a clinical setting. Performing a three-dimensional CFD analysis on a network of blood vessels has significant computation and time costs, making it sub-optimal for clinical use [10, 11]. This is the main reason as to why 3D CFD has not yet been employed to measure hemodynamic properties in large cohorts. However, there are alternative approaches.

This study will utilize computed tomography (CT) scans that are currently present in the PMBB. There have been previous studies that have utilized biobanks to gather a large cohort of subjects. One such study utilized magnetic resonance imaging (MRI) and neural networks to improve automatic segmentation of aortas in atypical pathologic cases. This study utilized MRI scans from the UK biobank, which allowed for a large and diverse dataset for the study [12]. For this study, the basic pipeline is that aortas will be segmented from CT scans contained in the PMBB, then geometric properties required for the 1D model will be extracted and input into the ROM with varying initial conditions for measurement of hemodynamic properties. While some information from a 3D CFD simulation may be lost, a ROM approach allows for significantly more simulations to be performed in the same amount of time, allowing for a whole characterization of an individual’s aortic flow response.

This paper utilizes a one-dimensional Windkessel model, which considerably reduces both the computational power and time required to perform CFD modeling, reducing the time required from hours to seconds [13]. Using this reduced order model (ROM) would then allow for a large-scale application of hemodynamic modeling of the aorta, which is the focus of this study. There has been previous work involving ROMs of blood vessels, however, these studies either have small datasets, making them only proofs of concepts, or they attempt to address the issue by generating synthetic datasets for their studies in the form of geometries and inflow data [14, 15]. This allows for a novel approach of modeling hemodynamic properties on a large scale using non-synthetic geometries gathered from the Penn Medicine Biobank (PMBB).

## Materials and Methods

### Penn Medicine Biobank

The PMBB recruits participants at Penn Medicine Health System (Philadelphia, PA) by enrolling them at the time of outpatient visits. Patients complete a questionnaire, donate a blood sample, and agree to future recontact. It is an ethnically diverse cohort with black patients composing of nearly 25% of participants. This study was approved by the Institutional Review Board of the University of Pennsylvania and all patients have given informed consent to participate in this study. All methods were performed in accordance with the relevant guidelines and regulations.

### Scan Acquisition and Preprocessing

From the PMBB, scans were selected from those being thoracic CT scans with the ability to have machine learning segmentation applied. The manufacturers of the CT scanners were Siemens, GE Medical Systems, Toshiba, and Siemens Healthineers, with slice thickness ranging from 0.5-1.5 mm. Each of these images were then segmented using TotalSegmentator to allow for collection of geometric parameters for the model [16]. Before acquiring geometric quantities from the segmentation, all aorta segmentations were clipped so that only the thoracic aorta was utilized. To select only the thoracic aorta, the segmentation was clipped at the center point of the T12 vertebrae. This location was chosen as the documented point where the descending thoracic aorta ends, and the abdominal aorta begins [17].

### Segmentations to Geometries

To apply the reduced order model to the segmentations, various geometric quantities had to be extracted from them. The program used to do this was the Vascular Modeling Toolkit (VMTK 1.4.0). First, we identified the aorta segmentation from the result of TotalSegmentator, and then generating a deformable model from this. An approximated centerline was then generated from this model, and then used to locate the source and target points which were next utilized for generating a more accurate centerline. This centerline in conjunction with the deformable model was then applied for generating various properties of the thoracic aorta including the length, average and max diameter, average and max curvature, and average and max torsion. To apply the diameter along the length of the thoracic aorta to the model, a polynomial fit of order 20 was applied to it, the resulting equation be used by the model. This was done in order to supply the model with a continuous equation, as that was what was required for it to run. The tapering angle of each ascending thoracic aorta was also calculated. The taper of a vessel is the rate at which the vessel reduces its diameter along its length. The tapering angle can be calculated from the equation below. Where D_max_ is the maximum diameter of the vessel, D_min_ is the minimum diameter of the vessel and L is the length of the vessel.

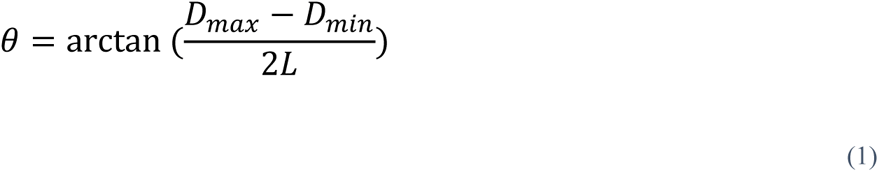

### Reduced Order Model Overview

The reduced order model is based on a 3 element Windkessel model, which is commonly used in the reduced order modeling of blood vessels, in conjunction with the fluid-structure interactions that occur from the distensibility of the aortic vessel wall [12]. This system requires three equations, mass conservation, momentum conservation and a relationship between pressure, area and distensibility of the vessel. These can be seen in equations 2, 3 and 4 respectively.

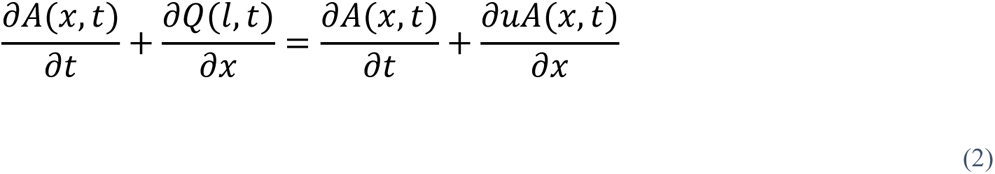

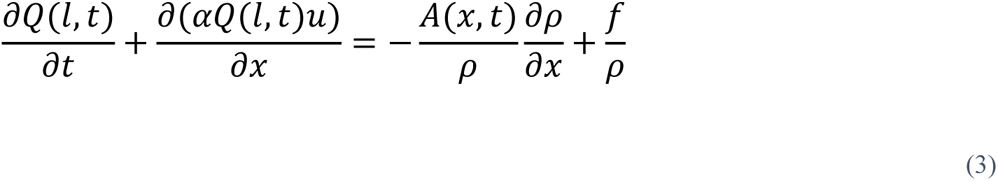

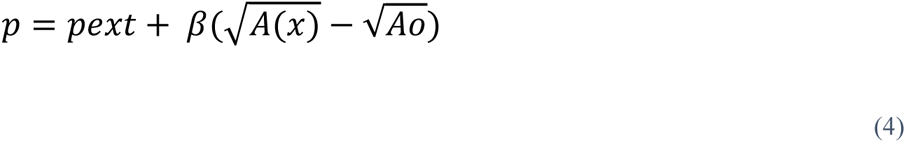

In these equations A describes the cross-sectional area of the vessel, Q describes the volumetric flow rate, α is a momentum flux correction factor set to 1.1 for parabolic flow, u is the flow velocity, ρ is the density of the fluid, *f* represents the force of friction in this system, *p* is the pressure inside the vessel *p_ext_* is the external pressure and *A_o_* is the area of the vessel at equilibrium. *β* is a term that is a property of the vessel and is calculated from equation 5.

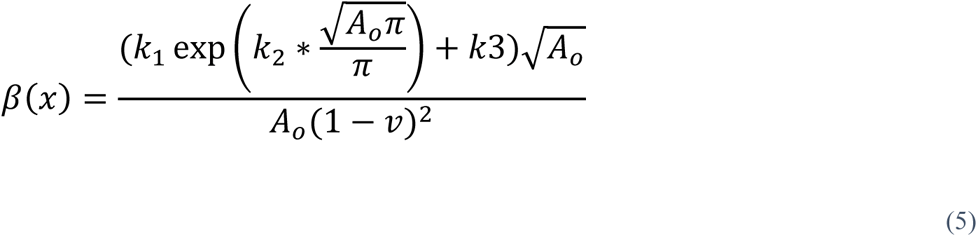

This beta term is described by *k_1_*, *k_2_*, and *k_3_*, which are constants that experimentally fit for calculating the ratio between the Young’s modulus and the dimensions of the vessel [18]. This system of equations describes the pulse wave propagation down the vessel. To transform the input waveform into a physiologically accurate waveform a 3-element Windkessel model is utilized. A Windkessel model describes fluid flow through a blood vessel in terms of an electronic circuit governed by equation 5 [19].

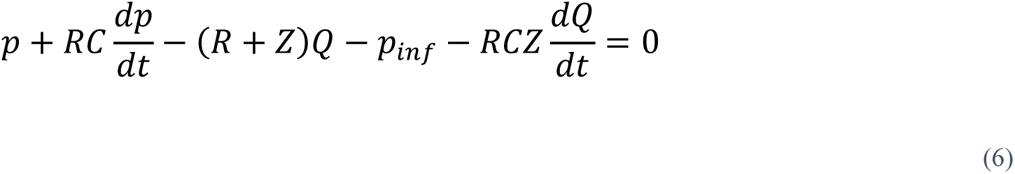

*R* being the peripheral resistance of the vessels, *C* being the vessel compliance and *Z* being the characteristic impedance of the vessel, which was calculated as from a Tau ratio, being the ratio of the peripheral resistance to the characteristic impedance (τ=R/Z), the Tau ratio was calculated from a previous study [20]. The tapering angle of each aorta was calculated by dividing the difference between the max and minimum aortic diameter by the length and taking the arctangent of that value.

### Parameter Variation

To generate the hemodynamic fingerprint, various parameters can be varied from the model. These include the peripheral resistance, vessel compliance and the input volumetric flow waveform. The peripheral resistance and vessel compliance values were gathered from previous literature discussing measurements of these through a range of patients [20–22]. The peripheral resistances varied between 710 dynes*s/cm^5^ to 2900 dynes*s/cm^5^, while the compliance remained constant. The basic flow used was a pulse sine wave that peaks at 200 cm^3^/s during systole and goes to 0 cm^3^ during diastole. This was varied by increasing and decreasing the peak flow rate by 10%, leading to a range of 180 cm^3^/s to 220 cm^3^/s. For the purposes of this study, the model was used by only varying the peripheral resistance and keeping the compliance at a constant value. The compliance was not varied to determine the effect of only parameter at a time. The entire pipeline from CT scan to ROM output can be seen in figure 1, the physical representation of the reduced order model can be found in figure 2. Simulated measurements were taken at the end of the aorta, at the T12 vertebrae.

**Figure 1.**
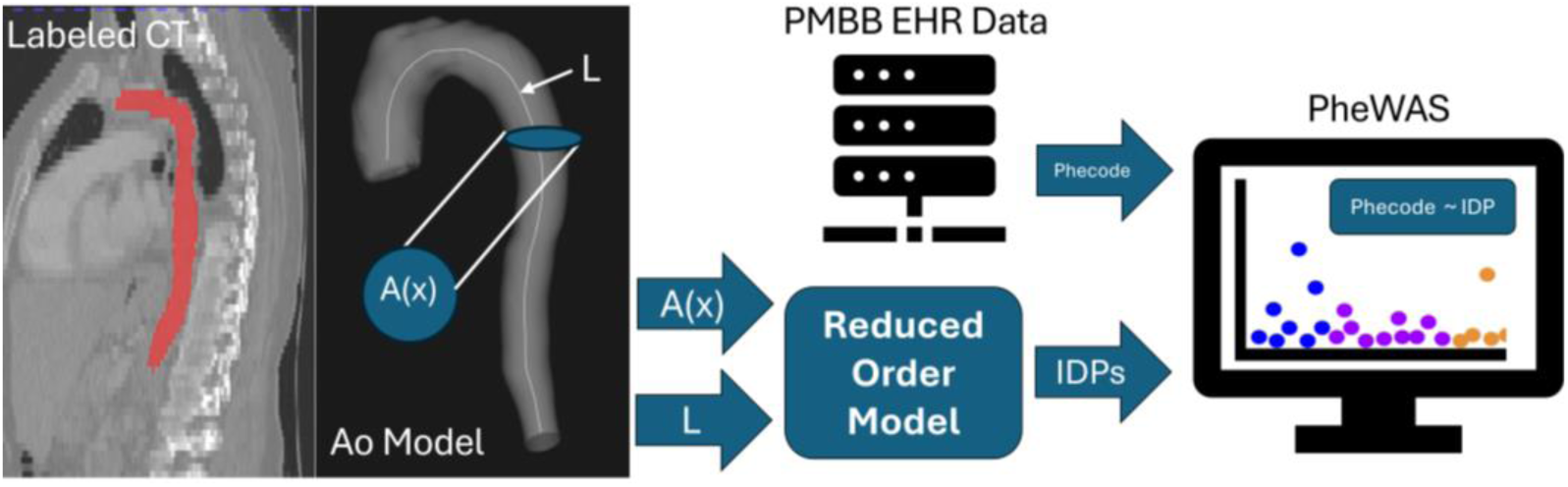
Processing pipeline from CT scan data in the PMBB to machine learning segmentation to geometry calculations and finally hemodynamic calculations from the ROM. Including an example VMTK geometry as well as the three element Windkessel model.

**Figure 2.**
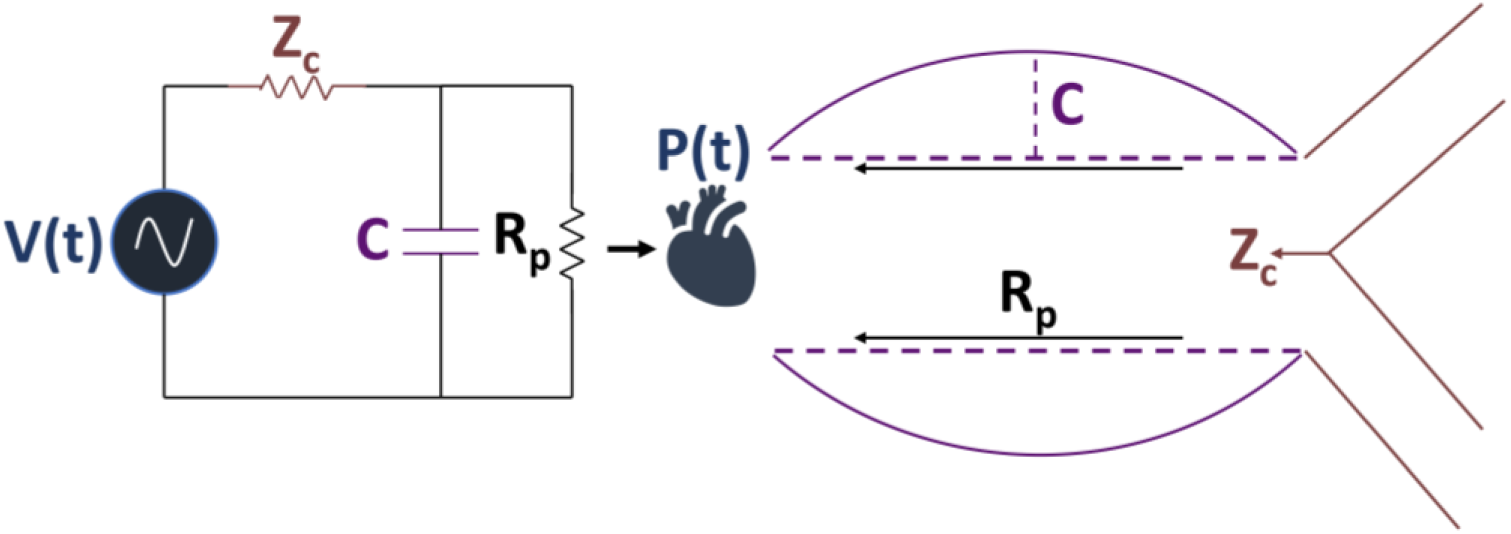
Representation of ROM in physical elements. V(t) corresponds to the pulsatile nature of the pressure waveform, Z_c_ represents the characteristic impedance of the vessel, C represents the compliance of the vessel and finally R_p_ is the peripheral resistance.

### Statistical Analysis

To examine the effects that geometric data has on different phecodes, a unique identifier for different pathological traits, a phenome wide association study (PheWAS) was applied. R Studio was used for all statistical analysis. This PheWAS consisted of a logistic regression of each phecode with multiple parameters for correction. We used a logistic regression to look at the association of the different phecodes with max aortic diameter, correction for thoracic aortic length, patient age, and patient sex. A second PheWAS was applied, this time looking at the results of the ROM. This logistic regression looked at the association between the different phecodes and the pulse pressure of the model divided by thoracic aortic length, correcting for patient age, patient sex, and max aortic diameter. Phecodes that had an occurrence of more than 100 times were plotted in each PheWAS.

## Results

### Thoracic aorta anatomic imaging traits and their association with disease

The investigators identified 3,216 thoracic CT scans from the Penn Medicine Biobank. After ML-derived aortic segmentation and ROM analysis, only 12 thoracic CTs were unable to be processed, resulting in a final data set of 3,204 thoracic CTs with imaging traits. The distribution of patient age among the CT scans is shown in Table 1, with most patients between 40 and 79 years old. To examine changes in aortic geometry, we investigated the aortic radius among participants by age (Figure 3A). The average thoracic aortic diameter was 26.6 ± 3.1 mm, and the average length was 310 ± 37 mm. (Figures 3B and 3C, respectively). Correlation between imaging traits is shown in Figure 3D. The most closely related imaging traits were aortic diameter and length. PheWAS associations between maximum aortic radius and disease are shown in Figure 4. The four phecodes that showed the highest association with aortic radius were aortic aneurysm (p = 2.29E-24), other aneurysms (p = 1.21E-18), nonrheumatic aortic valve disorders (p = 6.28E-13) and heart valve disorders (p = 8.22E-9) (Figure 3B).

**Figure 3.**
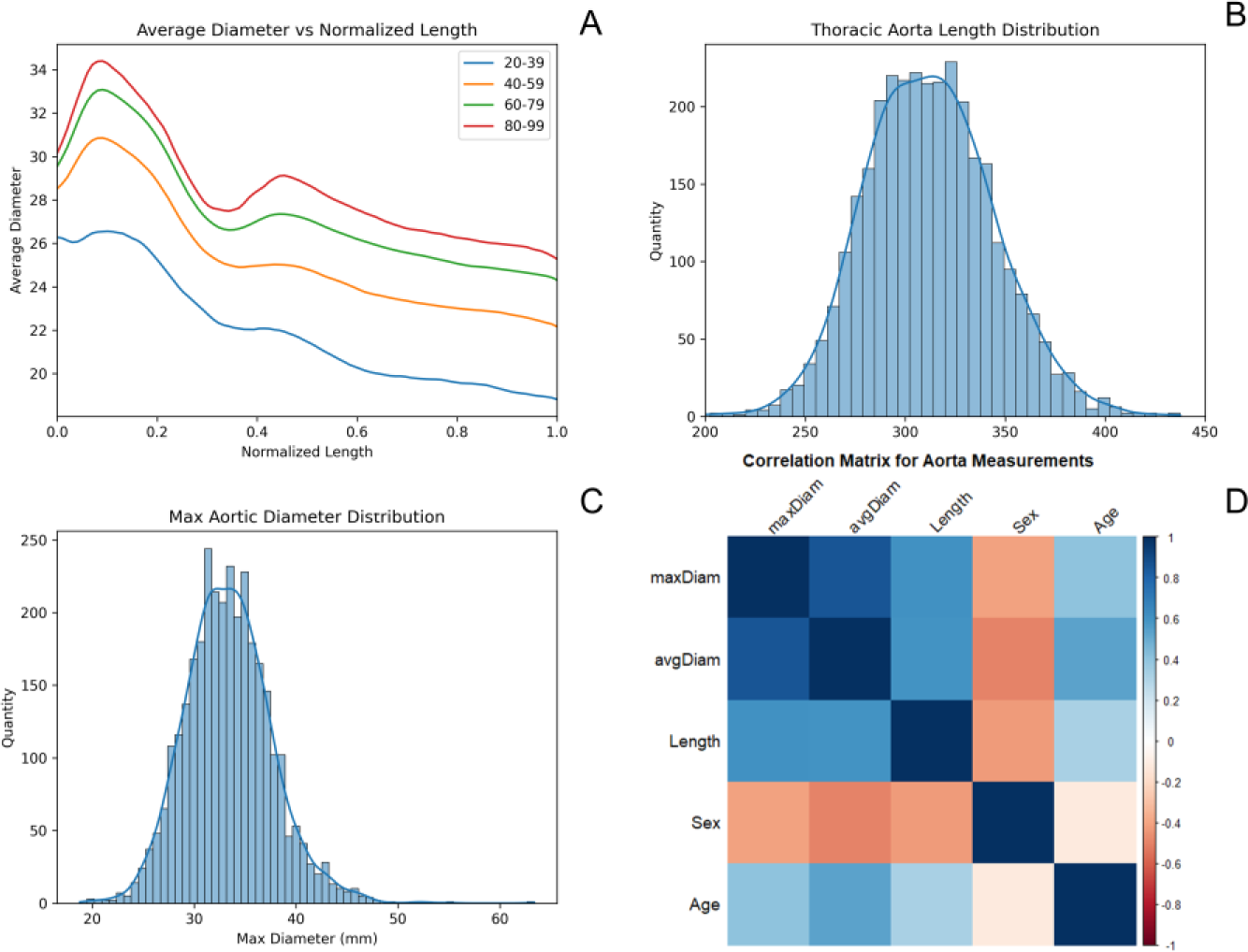
A) The average diameter versus the normalized length of each thoracic aorta for the different age groups (n = 3204). B) Max thoracic aorta diameter distribution (n = 3204). C) Thoracic aorta length distribution (n = 3204) D) Correlation graph between imaging traits between max thoracic aortic diameter, average thoracic aortic diameter, ascending thoracic aortic length, patient sex, and patient age.

**Figure 4.**
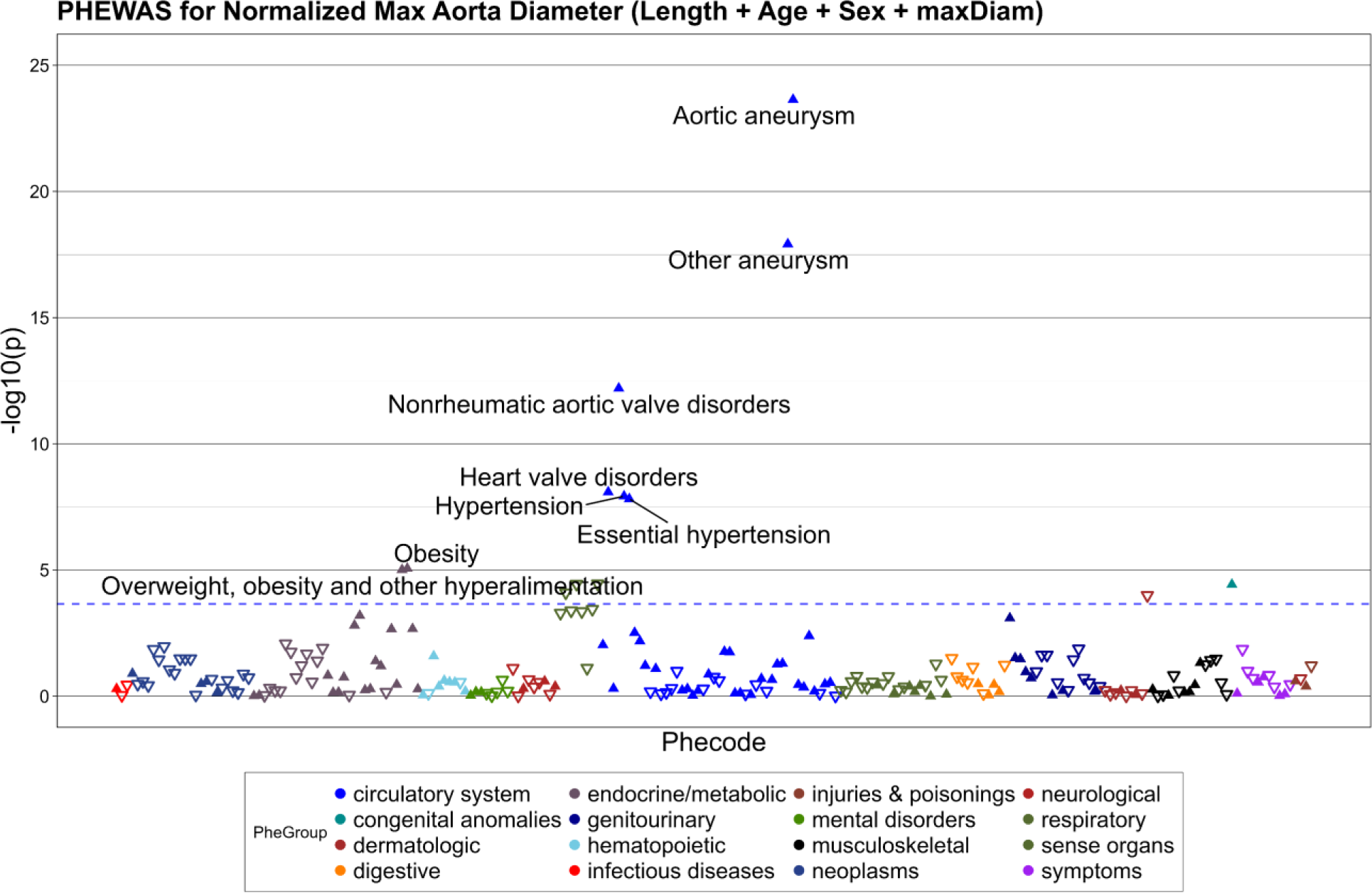
Phenome wide association study analyzing the effect max aortic diameter has on phecodes response (n = 1934)

**Table 1.** Description of the dataset including, age, sex and top 10 most prevalent phecodes for scans containing that information.

|  | N=3,204 |
| --- | --- |
| **Age** |  |
| <20-39 | 213 (6.6%) |
| 40-59 | 881 (27.5%) |
| 60-79 | 1914 (52.7%) |
| >79 | 196 (6.1%) |
| **Sex** |  |
| Male | 1617 (50.4%) |
| Female | 1587 (49.5%) |
| **Top 10 Phecodes** | N = 1934 |
| Hypertension | 1147 (59.3%) |
| Essential Hypertension | 1117 (57.8%) |
| Disorders of Lipid Metabolism | 1022 (52.8%) |
| Hyperlipidemia | 1021 (52.8%) |
| Symptoms of Respiratory System | 827 (42.8%) |
| Diseases of Esophagus | 767 (39.7%) |
| Esophagitis | 730 (37.7%) |
| GERD | 684 (35.4%) |
| Cardiac Dysrhythmias | 618 (32.0%) |
| Overweight/Obesity | 609 (31.5%) |

### ROM hemodynamic imaging traits and their association with disease

CFD simulations were performed with 8 different resistances, and 5 different inflow waveforms, resulting in over 125,000 simulations in 3,204 aortas. Figure 5 shows the time-varying aortic pressure for one cardiac cycle and its change with resistance. The peak pressure for the dataset was 201 mmHg, and the minimum pressure calculated by the model through the range of parameters was 63.6 mmHg. Figure 6A shows the pulse pressure with resistance and the input flow boundary conditions. Finally, figure 6B shows the pulse pressure distribution for the median resistance and flow boundary conditions. The average pulse pressure for this data set was 22.5 ± 8.5 mmHg.

**Figure 5.**
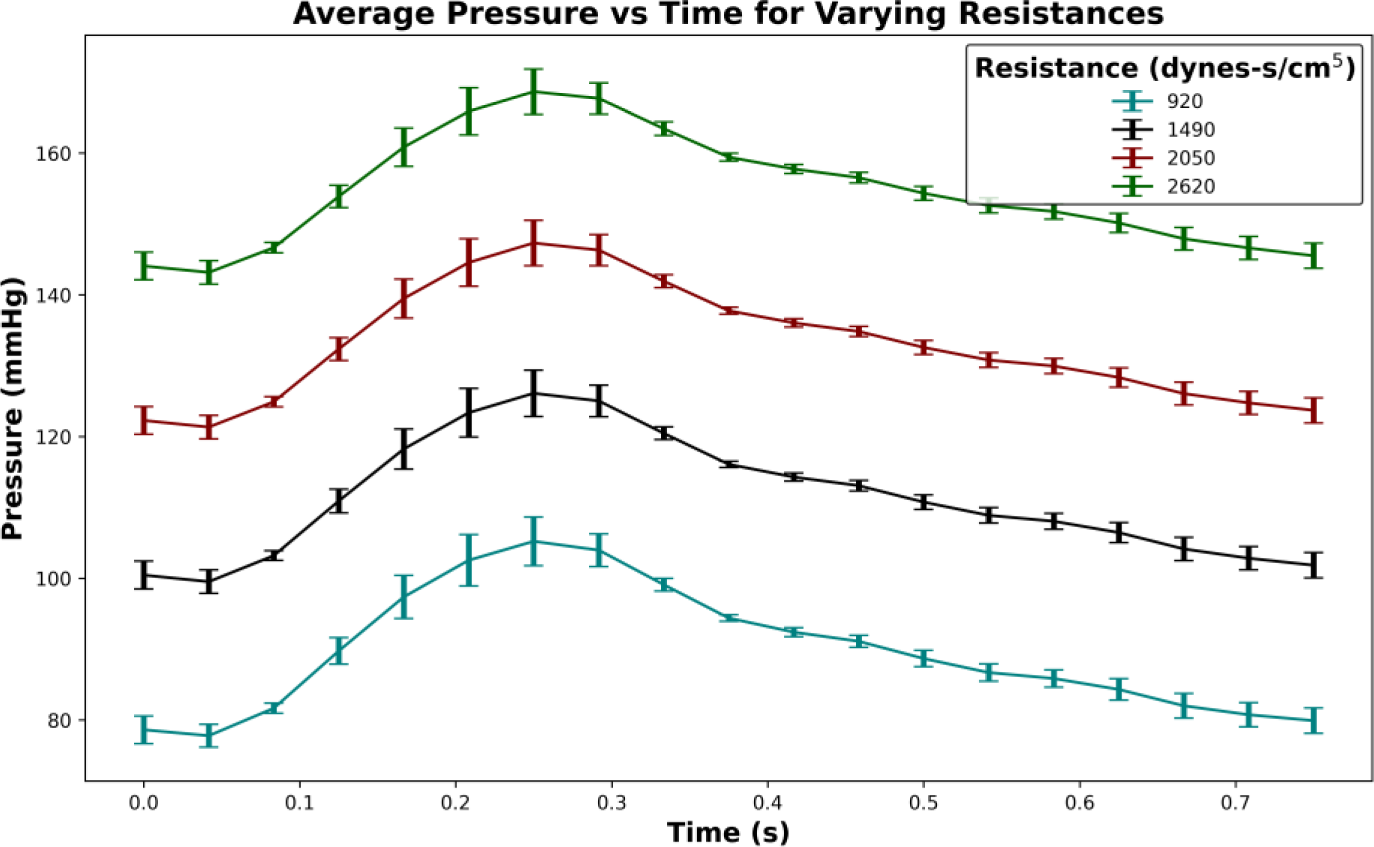
(n = 3204 for all) Pressure versus time curve for increasing resistances with a peak inflow value of 200 cm^3^/s. Showing variability between different geometries at each point. (n = 3204)

**Figure 6.**
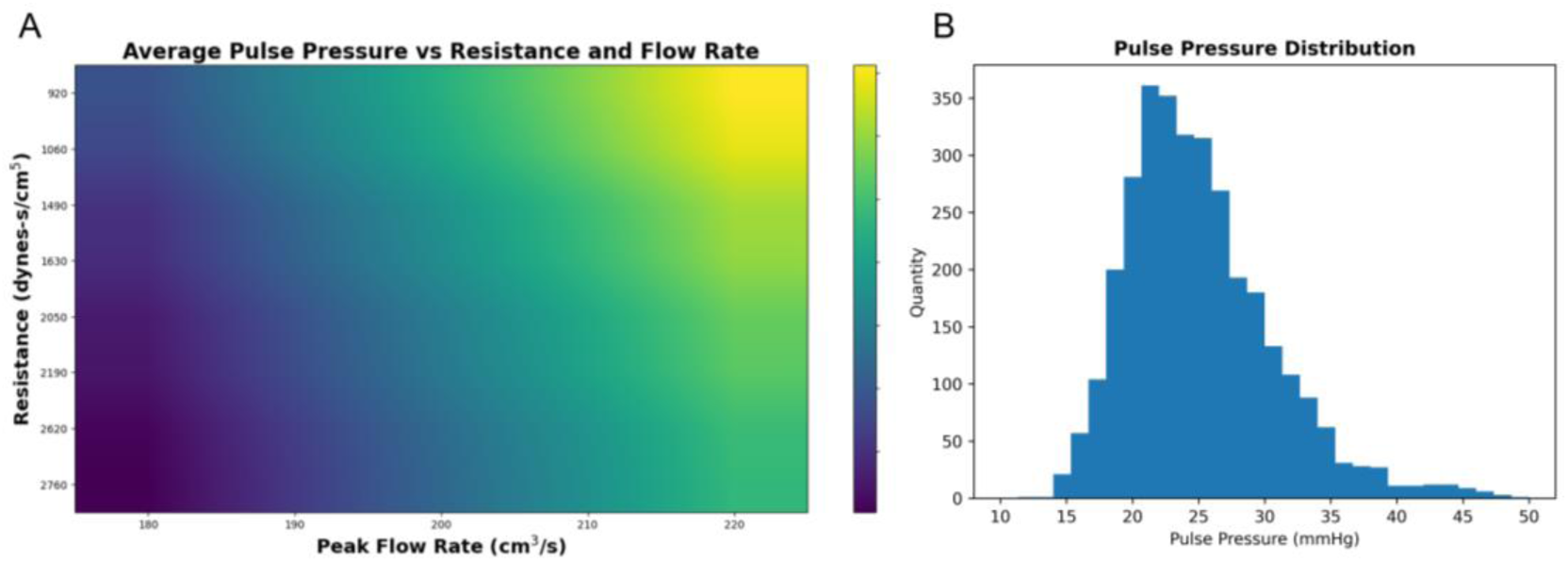
A) Surface plot showing how the average pulse pressure for each geometry changes with resistance and peak flow rate. B) Pulse Pressure distribution for all geometries for the median resistance and flow rate (n = 3204).

A PheWAS was performed between disease (phecodes) and aortic hemodynamic imaging trait pulse pressure, normalized by aortic length (Figure 7). The PheWAS showed that aortic aneurysm, nonrheumatic aortic heart valve disorders, heart valve disorders and other aneurysm were significantly associated with pulse pressure.

**Figure 7.**
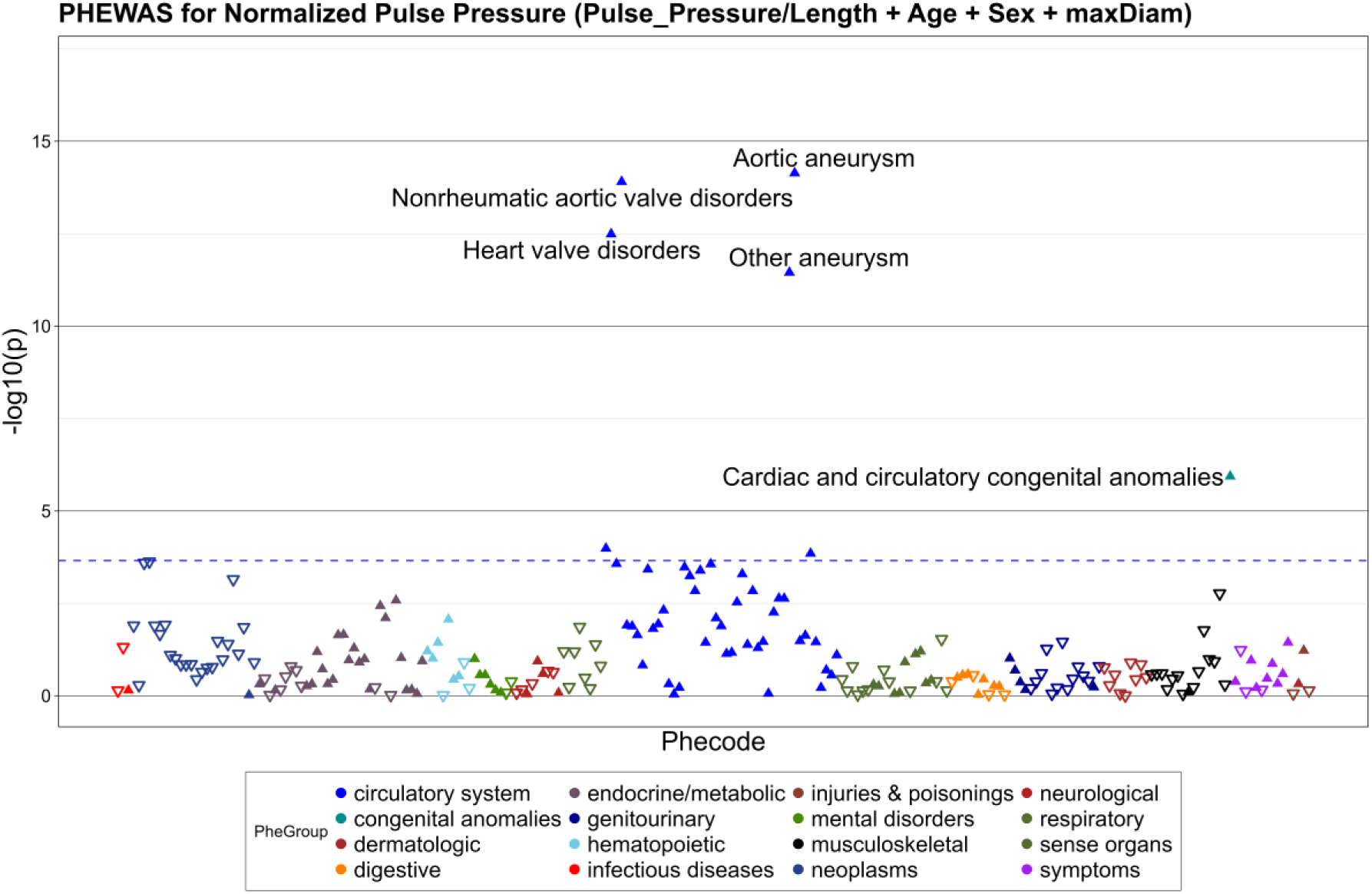
PHEWAS for Pulse Pressure normalized by thoracic aorta length, showing the most significant phecodes (n = 1934).

To further investigate the relationship between pulse pressure and significantly associated disease, the pulse pressure distribution, for a given resistance and flow rate, was examined by comparing patients with aortic aneurysm, other aneurysm, heart valve disorders, and aortic valve disorders to other PMBB patients (Figure 8). Patients with significantly associated aortic pathology showed a lower pulse pressure under the same boundary conditions.

**Figure 8.**
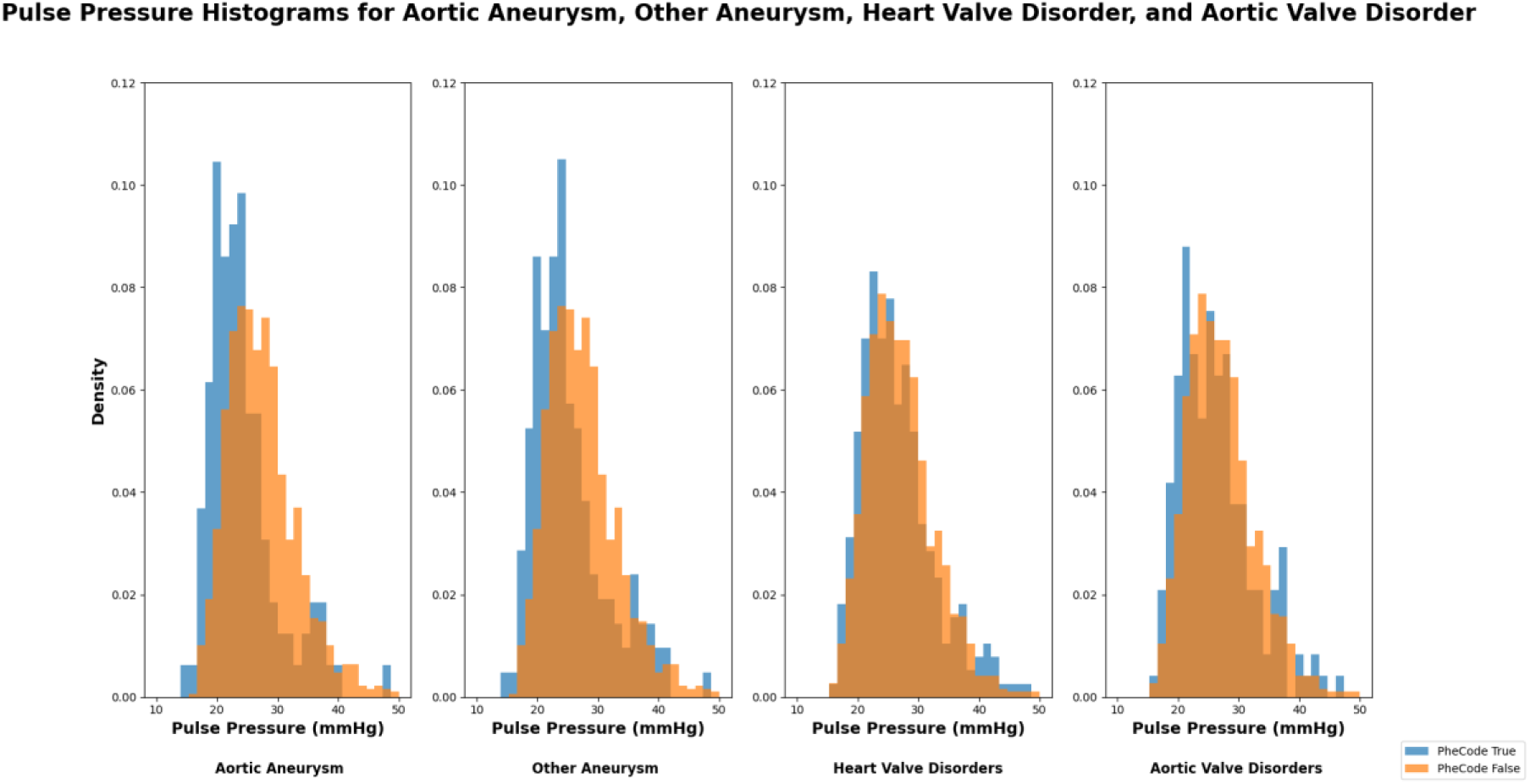
Example pulse pressure histogram for the four most significant phecodes resulting from the PheWAS. (aortic aneurysm n = 123, other aneurysm n = 158, heart valve disorders n = 293, aortic valve disorders n = 181)

To further examine how the geometry of the patient’s aorta affects the flow simulations, the tapering angle for the dataset was calculated for all 1934 patients present with pheCode data. The tapering angle is a parameter which characterizes the rate at which the aortic radius reduces over its length. A linear regression was then performed between the calculated pulse pressures and the tapering angle. There was a significant, but weakly negative correlation between the tapering angle and the pulse pressures calculated from the fluid simulations (p=0.048). The average tapering angle for this data set was 0.12 ± 0.04°.

## Discussion

This work provides a non-invasive approach of determining arterial hemodynamics from geometries gathered from medical imaging data. This was done at-scale by first calculating the geometries of 3,204 unique thoracic CT scans from a large-scale medical biobank using deep learning, then performing hemodynamic simulations across a range of wide range of clinically relevant hemodynamic parameters to generate a hemodynamic fingerprint of each aorta, when invasive measurements of flow and pressure are unknown. This proof-of-concept study shows the feasibility of applying ROMs to biobank patients at this scale of thousands of patients. An additional novel aspect of this work was to incorporate ROM hemodynamic analysis with clinical diagnostic data (phenome codes), showing that, when normalizing this calculated pulse pressure by the thoracic aorta length, specific, relevant phecodes for aortic disease are significantly associated with the derived hemodynamic parameters from the ROM.

While previous studies have incorporated patient-specific ROMs, these studies have not been applied on a large scale, as is possible with the PMBB [23]. There have also been large scale studies to examine geometric measurements such as coronary calcium and ventricular mass, to show how these lead to aortic disease [5]. However, these have not incorporated hemodynamic measurements into the studies. There have also been studies that aim to develop methodologies for quickly calculating CFD results using ROMs. A study by Biancolini et al. utilized 3D simulations calculated in ANSYS to develop a methodology for generating a digital twin for patient specific results [24]. This methodology is a similar approach to this study; however, this paper uses a variation of parameters was used to characterize an individual geometry. The UK Biobank has also been used to characterize aortic geometries previously, this was done to calculate parameters such as aortic distensibility, or genetic associations between different aortic valve diseases [25, 26]. By utilizing a ROM for flow simulations, we are able to quickly characterize an array of aortic geometries. By combining this approach with more classical examples of imaging characteristics, such as calcification of the aorta, novel observations can be made on a large scale.

By calculating the tapering angle of the ascending thoracic aortic dataset, a statistically significant but weakly negative correlation was determined between the calculated pulse pressure from the ROM and the tapering angle. Future work can examine further how the results of the reduce order model are related to the tapering angle, or other properties of thoracic aortic geometry.

This study attempted to examine the effects of aortic geometry on ROM hemodynamic measurements. There is potential for it to be used for other aortic diseases as well, for example in atherosclerosis. Atherosclerosis is an inflammatory disease that leads to plaque buildup on vasculature walls and often goes undiagnosed until a cardiac event occurs [24]. There is evidence that hemodynamic measurements like pulse wave velocity and pressure magnitude have a relevant impact on the formation of atherosclerotic plaques [9]. In the future, this work could be applied to examining the effects that ROM hemodynamic measurements have on formation of atherosclerotic plaque and could potentially supply an opportunistic screening method for early detection.

The major limitation of this study is the lack of inflow data due to using CT scans. If this methodology were incorporated for an imaging modality that does provide flow data, like 4D MRI, the process could become more patient specific by fitting the Windkessel parameters for that specific patient. In that case other parameters could be varied, such as the ones controlling the pulse wave propagation down the vessel. This study also only examined the simulated hemodynamics at a single point in the vessel. More work can be done by examining the pulse wave propagation as it progresses down the aorta. Another result of the ROM that can be examined in the future is pulse wave reflections, as there is evidence to them being clinically significant [25].

This work provided a methodology for gathering ROM hemodynamic information at a large scale and provided a basis for analyzing these results together with clinical information. This was done by first taking CT scan data from the PMBB, converting it to geometries that can be utilized by a ROM, and then simulating through a range of parameters. In the future work can be done to examine different outputs of the ROM, such as reflected waveforms and perform analysis on the clinical significance of them in the PMBB.

